# Sustained antigenic engagement programs multidimensional CTL states with distinct cytotoxic capacities

**DOI:** 10.64898/2026.10.12.758563

**Authors:** Hazyadee Frecia Rodriguez-Gutierrez, Ashwin Jainarayanan, Marie Tosolini, Fereshteh Aslani, Anne-Laure Iscache, Sabina Müller, Béatrice Corre, Alejandro Vanazzi, Mirudula Elanchezhian, Camille Laurent, Salvatore Valitutti, Roxana Khazen

## Abstract

Cytotoxic T lymphocytes (CTLs) eliminate infected or malignant cells through repeated target-cell encounters, yet how they sustain cytotoxic activity during extended antigenic stimulation remains poorly understood. In particular, it remains unclear whether preserved population-level killing reflects maintenance of a uniform effector phenotype or diversification into functionally distinct CTL populations. Here, we combined live-cell imaging, high-dimensional cytometry, single-cell RNA sequencing, phenotypic trajectory analysis, population sorting, and rechallenge killing assays to define how human clonal CTLs remodel during sustained antigenic engagement. Prior antigen encounter altered the kinetics and pattern of target-cell membrane permeabilization but did not compromise overall subsequent killing upon rechallenge. Single-cell transcriptomic profiling after prolonged engagement revealed broad remodeling of antigen-experienced CTLs and heterogeneous cytotoxic transcriptional programs. Time-resolved scRNA-seq further showed that these functional dimensions remodeled with distinct kinetics, with rapid induction of activation- and cytotoxicity-associated programs, later remodeling of vesicular-trafficking programs, and progressive reduction of adhesion/ contact interaction-associated programs. High-dimensional cytometry showed that this temporal remodeling was accompanied by progressive redistribution of CTLs across a multidimensional protein-expression landscape defined by distinct combinations of cytotoxic, vesicular/degranulation, activation, cytokine, adhesion, and regulatory features. Phenotypic trajectory analysis organized this protein landscape along a temporal remodeling axis, with early lytic-granule-rich profiles giving way to multiple late-engaged phenotypes with distinct combinations of functional protein features. Surface-marker-defined late CTL populations displayed distinct killing capacities upon rechallenge, demonstrating that preserved population-level cytotoxicity can coexist with substantial functional heterogeneity. Finally, CAR T cells exposed to CD19^+^ targets retained cytotoxic activity upon rechallenge and acquired multidimensional phenotypic features that partially overlapped with the remodeling framework identified in clonal CTLs. Reanalysis of longitudinal patient-derived CAR T-cell transcriptomes further showed that related functional programs were dynamically organized after infusion, with the features distinguishing responder and non-responder tisa-cel samples changing between the infusion product and day 7. Together, these findings indicate that sustained CTL killing is accompanied by progressive multidimensional remodeling and diversification rather than maintenance of a fixed effector phenotype. Our findings support a model in which population-level cytotoxic activity can be maintained while the CTLs composing that population progressively diversify into functionally distinct states during sustained antigenic engagement.

## Introduction

CD8⁺ cytotoxic T lymphocytes (CTLs) are central effectors of immune responses against infected and malignant cells. Upon recognition of cognate peptide-MHC class I complexes, CTLs form a lytic synapse that supports polarized delivery of perforin, granzymes, death-receptor ligands, cytokines, and supramolecular attack particles(Bálint et al., 2020; Cassioli & Baldari, 2022; Chang et al., 2022; Filali et al., 2022; McKenzie et al., 2022). These mechanisms allow CTLs to trigger rapid and spatially confined target-cell death through multiple molecular routes, including apoptotic and non-apoptotic programs (McKenzie et al., 2022). CTLs are also potent serial killers. Individual cells can engage and eliminate multiple targets sequentially and, in some settings, attack several target cells within overlapping time windows, allowing relatively small effector populations to control large numbers of susceptible targets (Dekkers et al., 2023; Ronteix et al., 2022; Weigelin et al., 2021; Wiedemann et al., 2006). Such repeated cycles of target-cell recognition, synapse formation, cytotoxic execution and disengagement impose a substantial functional demand on CTLs, requiring continuous coordination of target-cell scanning, lytic machinery deployment, synapse termination and readiness for subsequent encounters.

CTL cytotoxicity is heterogeneous even within clonally derived populations expressing the same antigen receptor (Cazaux et al., 2019; Halle et al., 2016; Khazen et al., 2021; Varadarajan et al., 2012; Vasconcelos et al., 2015). Individual CTLs differ in the timing and efficiency of killing, their capacity to eliminate multiple targets, and the balance between cytolytic activity and cytokine production. Only a fraction of CTLs may execute rapid or serial killing, whereas others induce delayed, incomplete, or non-lethal target-cell responses(Cazaux et al., 2019; Khazen et al., 2021; Vasconcelos et al., 2015). Thus, clonality and shared antigen specificity do not ensure functional uniformity. Rapid lytic events can also be uncoupled from longer stimulatory interactions, indicating that cytotoxicity, activation, cytokine production, and sustained contact are not necessarily synchronized(Faroudi et al., 2003; Purbhoo et al., 2004).

This heterogeneity raises an important question regarding how CTL functional phenotypes are defined. Cytotoxic capacity is often inferred from individual markers, such as granzyme B, IFNγ, CD107a, or selected activation receptors. However, target-cell killing depends on multiple cellular components, including cytotoxic cargo, vesicular and degranulation machinery, adhesion and contact-interaction molecules, cytokines, and activation or regulatory receptors. CTL function may therefore be better represented by combinations of these features than by any single marker.

Although CTL differentiation, exhaustion, and cytotoxic modalities have been extensively studied, less is known about how these functional dimensions are reorganized during the early hours of sustained target-cell engagement. In particular, it remains unclear whether cytotoxic, vesicular, activation, contact-interaction, cytokine, and regulatory programs remodel with similar kinetics, and how their reorganization relates to subsequent killing capacity. The central unresolved question is whether sustained population-level cytotoxicity reflects maintenance of a common effector state or diversification into phenotypically and functionally distinct CTL populations.

This question is also relevant to CAR T-cell therapy. Durable tumor control requires CAR T cells to remain functional during repeated tumor-cell encounters, yet clinical outcome is influenced by differentiation state, product composition, exhaustion-associated programs, and functional heterogeneity(Haradhvala et al., 2022; Romain et al., 2022). Whether repeated tumor-cell engagement reshapes CAR T-cell heterogeneity through coordinated changes in cytotoxic, activation, adhesion and regulatory programs remains poorly understood.

Here, we combined live-cell imaging, high-dimensional flow cytometry, single-cell RNA sequencing, phenotypic trajectory analysis, population sorting, and functional rechallenge assays to address how functional heterogeneity develops within clonal CTLs during sustained antigenic engagement and whether the resulting phenotypes differ in their capacity to kill again. We show that prior antigen encounter alters target-cell death kinetics without compromising subsequent population-level killing. Sustained engagement redistributes CTLs across multidimensional protein and transcriptional landscapes characterized by different combinations of cytotoxic, vesicular, activation, contact-interaction, cytokine, and regulatory features. Time-resolved analysis further reveals that these functional dimensions remodel with distinct kinetics. Functional isolation of late-engaged populations demonstrates that the resulting phenotypes possess different killing capacities upon rechallenge. Finally, CAR T cells acquire partially overlapping remodeling features during repeated lymphoma-cell engagement. Together, these findings indicate that sustained cytotoxic activity is accompanied by progressive diversification of the CTL population rather than maintenance of a uniform effector phenotype.

## Results

### Prior antigen encounter alters target-cell death kinetics without compromising subsequent CTL killing

To determine whether CTLs retain cytotoxic function after prior antigenic engagement, we compared CMV-specific clonal CTLs during their first encounter with pp65 peptide-pulsed JY target cells with CTLs that had been pre-exposed to peptide-pulsed targets for 4 h and were then rechallenged with a fresh batch of target cells. Target-cell death was monitored by time-lapse microscopy in the presence of propidium iodide (PI), allowing us to follow both the timing and pattern of membrane permeabilization during CTL-mediated killing.

Non-exposed CTLs induced rapid and extensive PI uptake in target cells, whereas pre-exposed CTLs showed a more heterogeneous and overall delayed pattern of PI accumulation upon rechallenge (Fig. 1A-B). Consistent with this altered death dynamic, the time required for target cells to develop extensive membrane blebbing was significantly increased after conjugation with pre-exposed CTLs compared with non-exposed CTLs (Fig. 1C). Flow-cytometric analysis further showed that target cells engaged by pre-exposed CTLs displayed reduced PI uptake intensity, with a particularly marked reduction in the intermediate PI-positive population compared with targets engaged by non-exposed CTLs, a pattern consistent with reduced perforin-mediated membrane damage (Fig. 1D). Thus, prior cytotoxic engagement altered both the kinetics and quantitative pattern of target-cell membrane permeabilization.

**Figure 1.**
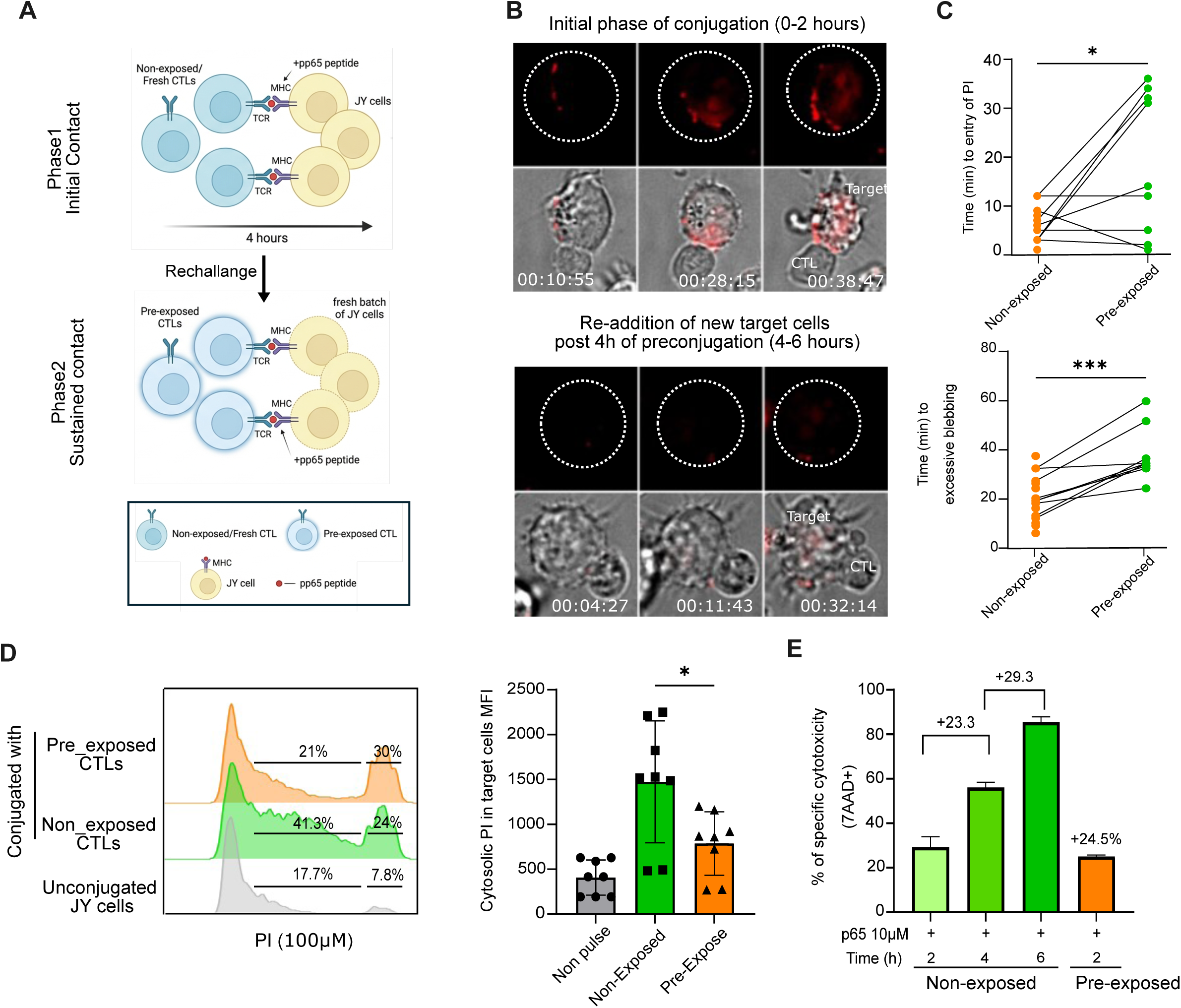
Prior antigen encounter alters target-cell death kinetics without compromising subsequent CTL killing. (A) Experimental design. CMV-specific clonal CTLs were conjugated with pp65 peptide-pulsed JY cells for 4 h, then rechallenged by adding fresh peptide-pulsed targets. (B) Representative time-lapse microscopy images during the first target-cell encounter (non-exposed CTLs, upper panel) or upon rechallenge after 4 h of prior engagement (pre-exposed CTLs, lower panel). Target-cell death was monitored by propidium iodide (PI) uptake over 2 h of live imaging. (C) Quantification of the time required for initial PI entry (upper panel) into target cells after conjugation with non-exposed or pre-exposed CTLs. Each point represents one CTL/target conjugate; n = 10 conjugates per condition. Quantification of the time to extensive membrane blebbing in target cells (lower panel) after conjugation with non-exposed or pre-exposed CTLs. Each point represents one CTL/target conjugate; n = 10 conjugates per condition. Data are from three independent experiments. (D) Flow-cytometric analysis of PI uptake in target cells following conjugation with non-exposed or pre-exposed CTLs. Representative histograms are shown on the left. The percentage of intermediate PI-positive population indicates differences in PI uptake intensity between target cells killed by non-exposed and pre-exposed CTLs are indicated on the histograms. Pooled quantification of mean fluorescent intensity (MFI) of PI is shown on the right; n = 3 independent experiments. (E) Quantification of CTL-mediated target-cell killing over 2 h in non-exposed and pre-exposed conditions. Pooled data are shown from at least three independent experiments. Data are shown as mean +/-SEM. Statistical significance was assessed using an unpaired Student’s t-test, where indicated.

Despite these changes in target-cell death dynamics, prior antigen encounter did not compromise subsequent CTL killing upon rechallenge with fresh peptide-pulsed target cells. Quantification of target-cell killing over the 2 h assay window showed that pre-exposed CTLs retained comparable overall cytotoxic activity under these conditions (Fig. 1E).

Together, these findings indicate that prior antigen encounter modifies the dynamics of CTL-mediated target-cell killing without measurably reducing overall cytotoxic capacity under rechallenge conditions. This observation raised the possibility that sustained antigenic engagement is accompanied by phenotypic remodeling rather than loss of cytotoxic function.

### Prolonged antigenic engagement reorganizes the CTL transcriptomic landscape and reveals heterogeneous cytotoxic transcriptional programs

Having found that prior antigen encounter altered target-cell death dynamics without compromising overall cytotoxic capacity, we next asked how sustained antigenic engagement reorganizes CTLs at the transcriptional level. We therefore performed single-cell RNA sequencing of CMV-specific clonal CTLs cultured for 16 h either alone, with unpulsed JY target cells, or with pp65 peptide-pulsed JY target cells. Integration of the three experimental conditions revealed a shared transcriptional landscape in which CTLs exposed to antigen-bearing target cells showed a markedly altered distribution compared with CTLs cultured alone or with unpulsed target cells (Fig. 2A).

**Figure 2.**
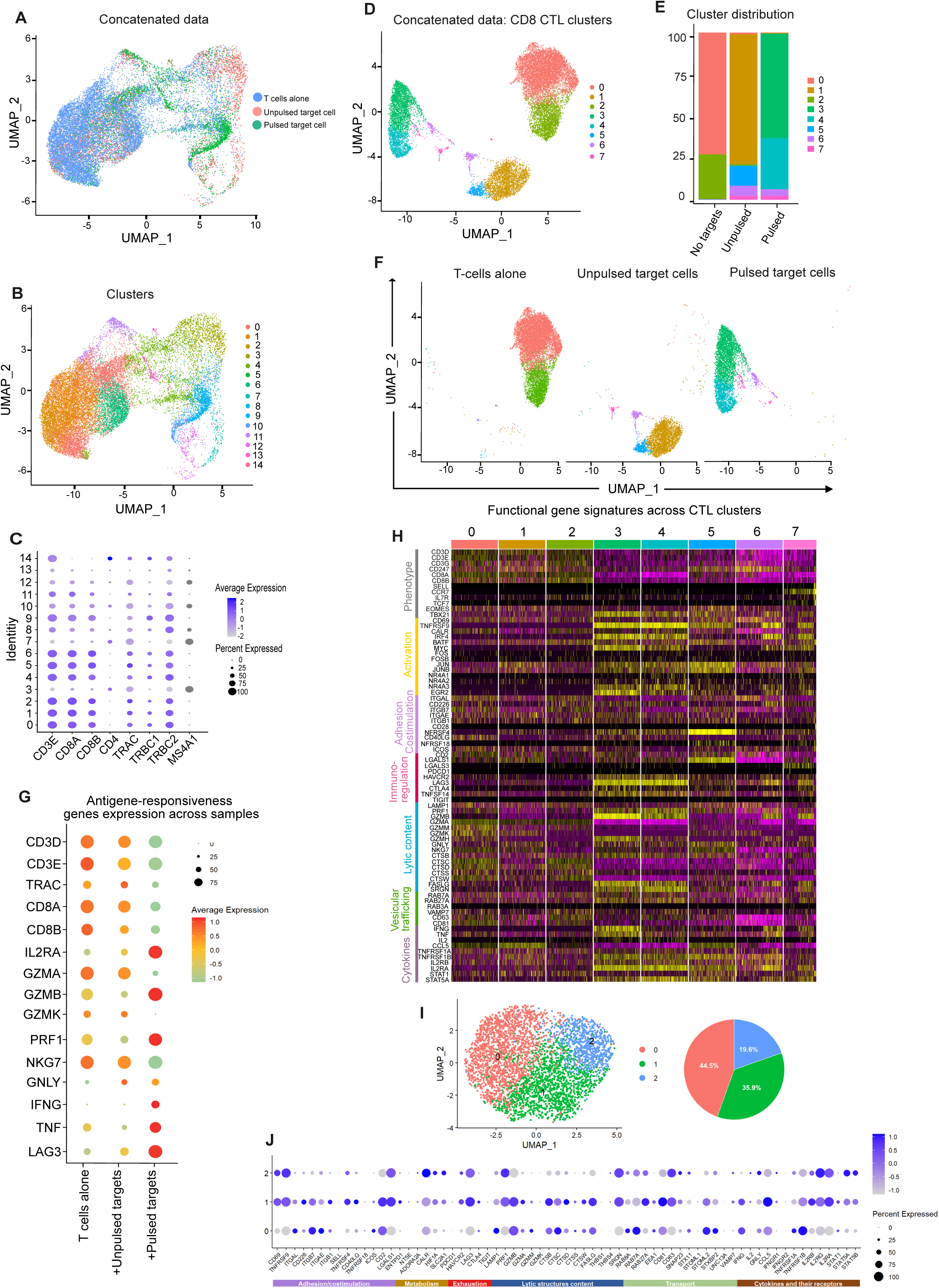
Prolonged antigenic engagement reorganizes the CTL transcriptomic landscape. (A) UMAP projection of single-cell transcriptomes from CMV-specific clonal CTLs cultured for 16 h either alone (blue), with unpulsed target cells (red), or with pp65 peptide-pulsed target cells (green). Cells are colored according to experimental condition to show the distribution of each sample within the integrated transcriptional landscape. (B) UMAP projection of the concatenated dataset showing unsupervised cell clusters identified across all experimental conditions. (C) Dot plot showing expression of CD8 T cell-associated genes across clusters, used to identify and select CTL clusters for downstream analysis. (D) UMAP projection of selected CD8 T cell clusters after exclusion of non-CTL populations. (E) Quantification of the relative distribution of selected CD8 T cell clusters across the three experimental conditions. (F) UMAP visualization highlighting the distribution of CTLs from each experimental condition within the selected CD8 T cell transcriptional landscape. (G) Dot plot showing expression of antigen-responsiveness genes across CTLs cultured alone, with unpulsed target cells, or with pp65 peptide-pulsed target cells. Dot size indicates the percentage of cells expressing each gene, and color indicates average expression. (H) Heatmap showing expression of selected functional gene signatures across selected CD8 T cell clusters. Genes are grouped according to T cell phenotype, activation/contact-associated phenotype, adhesion, regulatory/inhibitory receptor-associated programs, cytotoxicity, vesicular trafficking, and cytokine-associated categories. (I) Subclustering of antigen-engaged CTLs within the cytotoxic transcriptional region identified after 16 h conjugation with pp65 peptide-pulsed target cells. The UMAP shows three transcriptionally distinct subclusters, and the pie chart indicates their relative abundance. (J) Dot plot showing expression of selected functional genes across the antigen-engaged CTL subclusters shown in panel I. Genes are grouped by activation/regulatory, metabolic, cytotoxic, lytic-granule-content, trafficking, and cytokine/cytokine-receptor-associated categories. Dot size indicates the percentage of cells expressing each gene, and color indicates average expression. Data are from single-cell RNA sequencing of CMV-specific clonal CTLs after 16 h culture under the indicated conditions.

Unsupervised analysis partitioned the integrated dataset into transcriptionally distinct cell clusters (Fig. 2B). Expression of CD8 T cell-associated genes was used to identify CTL clusters and exclude non-CTL populations from subsequent analyses (Fig. 2C-D). Quantification of the selected clusters across experimental conditions revealed substantial differences in their relative representation, with antigen-engaged CTLs preferentially populating specific regions of the transcriptional landscape (Fig. 2E-F). Thus, prolonged recognition of antigen-bearing target cells was associated with a broad redistribution of clonal CTLs across transcriptionally defined populations.

We next examined the molecular programs underlying this redistribution. CTLs cultured with peptide-pulsed target cells exhibited increased expression of antigen-responsiveness genes compared with CTLs cultured alone or with unpulsed targets (Fig. 2G). Analysis of selected functional gene signatures further showed that the transcriptional clusters differed across multiple dimensions, including T cell phenotype, activation and contact-associated features, adhesion, regulatory and inhibitory receptor-associated programs, cytotoxicity, vesicular trafficking, and cytokine-associated genes (Fig. 2H). The clusters were therefore distinguished not by isolated changes in individual genes, but by different combinations of transcriptional programs spanning several functional dimensions of the CTL phenotype.

To further resolve heterogeneity within the antigen-engaged population, we subclustered CTLs located within the cytotoxic transcriptional region after 16 h of conjugation with peptide-pulsed target cells. This analysis identified three transcriptionally distinct subclusters with different relative abundances (Fig. 2I). The subclusters exhibited differential combinations of activation and immune-regulatory genes, cytotoxic machinery, lytic-granule components, trafficking-associated genes, and cytokine-or cytokine-receptor-associated genes (Fig. 2J). These findings reveal substantial transcriptional heterogeneity even among antigen-engaged CTLs occupying a broadly cytotoxic transcriptional region.

Together, these data show that prolonged encounter with antigen-bearing target cells is associated with broad transcriptional remodeling of clonal CTLs and reveals heterogeneous cytotoxic transcriptional programs within the antigen-engaged population. This heterogeneity prompted us to next investigate how the different functional components of the CTL transcriptional response develop over time.

### Time-resolved transcriptomics reveals rapid cytotoxic activation followed by delayed vesicular and contact-interaction remodeling

Having identified broad transcriptional remodeling and heterogeneity among antigen-engaged CTLs after 16 h, we next asked how the different functional components of this response developed over time. We therefore performed time-resolved single-cell RNA sequencing of CMV-specific clonal CTLs analyzed alone or after 2 h, 4 h, 6 h, or 16 h of conjugation with pp65 peptide-pulsed JY target cells.

Integration of all time points and unsupervised clustering revealed a time-dependent redistribution of CTLs across the transcriptional landscape (Fig. 3A). Although cells from different time points occupied partially overlapping regions, their relative distribution changed progressively during antigenic engagement. Visualization of clusters enrichment per sample confirmed that individual transcriptional clusters were differentially represented at early, intermediate, and late time points (Fig. 3B), indicating that sustained engagement was associated with progressive changes in the composition of the CTL population.

**Figure 3.**
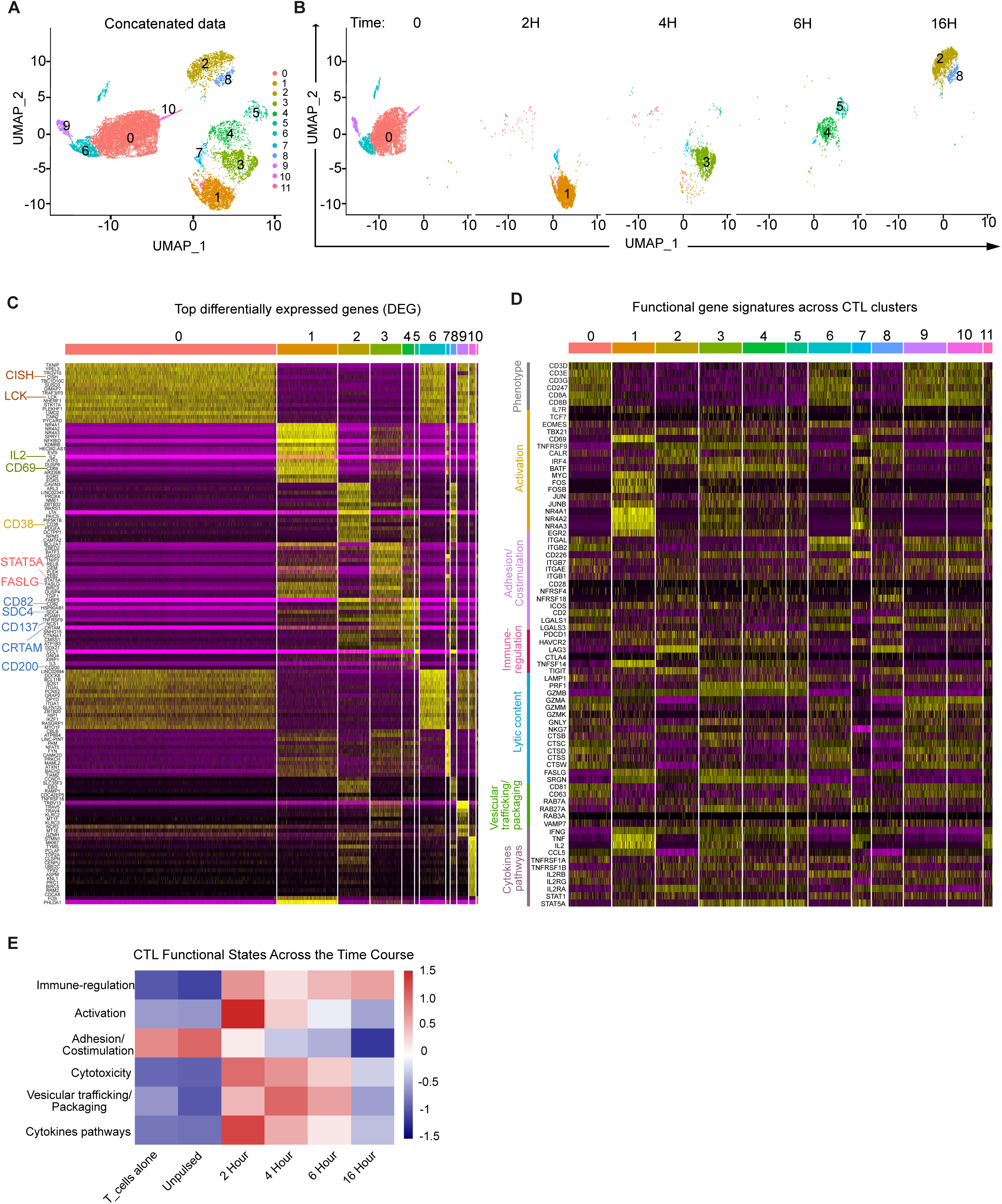
Time-resolved scRNA-seq reveals rapid cytotoxic activation followed by delayed vesicular and contact-interaction remodeling. (A) UMAP projection of single-cell transcriptomes from CMV-specific clonal CTLs analyzed across a time course of antigenic target-cell engagement. CTLs were analyzed alone, or after 2 h, 4 h, 6 h, or 16 h conjugation with pp65 peptide-pulsed JY target cells. Cells are colored by unsupervised cluster identity. (B) UMAP projections highlighting the distribution of CTLs from each experimental time point within the integrated transcriptional landscape. (C) Differentially expressed genes across CD8 T cell clusters, showing transcriptional programs associated with each condition. (D) Heatmap showing expression of selected functional gene signatures across transcriptional clusters. Genes are grouped according to T cell phenotype, activation/contact-associated phenotype, adhesion/contact interaction, regulatory/inhibitory receptor-associated programs, cytotoxicity, vesicular trafficking, and cytokine-associated categories. (E) Module scores for selected functional gene programs, including cytotoxic phenotype, activation, adhesion/contact interaction, vesicular trafficking, cytokine-associated genes, and regulatory/inhibitory receptor-associated genes. Data are from single-cell RNA sequencing of CMV-specific clonal CTLs isolated after the indicated durations of antigenic target-cell engagement. Module scores were calculated using predefined gene sets for each functional category.

Analysis of differentially expressed genes identified distinct transcriptional programs across the CTL clusters and experimental time points (Fig. 3C). Early antigen engagement was associated with increased expression of immediate activation- and cytotoxicity-associated genes, whereas later time points showed broader reorganization of vesicular-trafficking, adhesion/ contact interaction, and regulatory/inhibitory receptor-associated features. Analysis of selected functional gene signatures further demonstrated that the transcriptional clusters were distinguished by different combinations of T cell phenotype, activation/contact-associated, adhesion/ contact interaction, cytotoxicity, vesicular trafficking, cytokine-associated, and regulatory/inhibitory receptor-associated programs (Fig. 3D). Thus, the multidimensional CTL response did not develop as a single uniform program, but through temporally distinct changes across several functional components.

Module-score analysis further resolved the temporal dynamics of these transcriptional programs (Fig. 3E). Functional programs associated with cytotoxic effector molecules and cytokine production increased rapidly at early time points, in parallel with activation-related scores, whereas adhesion/ contact interaction-associated programs progressively declined during antigenic engagement. Programs related to vesicular trafficking and granule packaging peaked at intermediate time points, following the increase in expression of cytotoxic effector molecules. Regulatory and inhibitory receptor-associated programs also evolved dynamically over the time course. Together, these distinct kinetics indicate that the different functional dimensions of the CTL response are remodeled asynchronously rather than as part of a uniformly increasing program.

Together, these data reveal a temporally ordered and partially asynchronous remodeling of the CTL transcriptional response during sustained antigenic engagement, characterized by rapid induction of cytotoxic and cytokine-associated programs, intermediate remodeling of vesicular-trafficking and granule-packaging programs, and a progressive decline in adhesion/ contact interaction-associated programs. We next asked whether this transcriptional reorganization was accompanied by corresponding changes in the CTL protein-expression landscape.

### Multidimensional protein-expression patterns define CTL phenotypic remodeling during sustained antigenic engagement

Having identified temporally distinct transcriptional programs during sustained antigenic engagement, we next asked how these changes were reflected at the protein level and whether CTLs similarly diversified across multidimensional protein phenotypes. Because CTL function is often inferred from the expression of individual markers, we sought to determine whether prolonged antigenic stimulation instead reorganizes CTLs across multidimensional protein phenotypes. To address this, we performed high-dimensional flow cytometry on CMV-specific clonal CTLs analyzed alone or after 2 h, 4 h, or 16 h of conjugation with pp65 peptide-pulsed JY target cells.

Single-cell t-SNE analysis revealed a progressive redistribution of CTLs across the protein-expression landscape over time. CTLs cultured alone occupied defined regions of the t-SNE space, whereas antigen-engaged CTLs progressively redistributed across partially distinct regions following 2 h, 4 h, and 16 h of antigenic engagement (Fig. 4A). Feature plots demonstrated heterogeneous expression of proteins associated with activation, cytotoxicity, vesicular trafficking and degranulation, cytokine production, and inhibitory receptor expression across the landscape, showing that the observed redistribution involved variation across multiple protein features rather than changes in any single marker (Fig. 4B).

**Figure 4.**
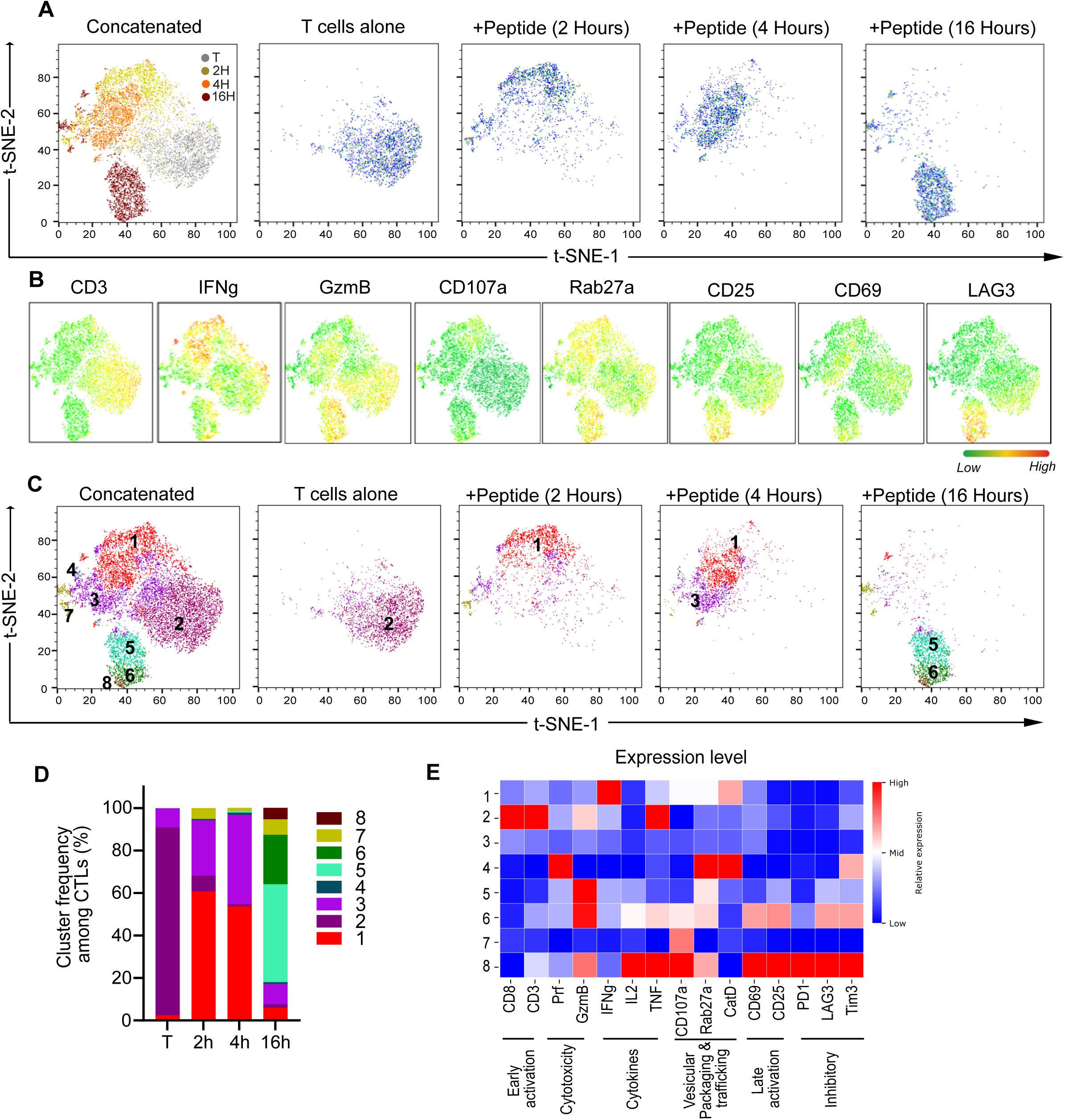
Time-dependent remodeling of the CTL protein-expression landscape during sustained antigenic engagement. (A) t-SNE projection of high-dimensional flow cytometry profiles from CMV-specific clonal CTLs analyzed alone or after 2 h, 4 h, or 16 h of conjugation with pp65 peptide-pulsed JY target cells. The concatenated map shows the integrated protein-expression landscape colored by experimental time point. Individual maps show the distribution of CTLs from each condition within the same t-SNE space. (B) t-SNE feature plots showing the expression of selected markers across the concatenated protein-expression landscape. Color scale indicates relative expression intensity from low (green) to high (red). (C) Protein-defined clustering of CTLs within the t-SNE landscape. The concatenated map shows all identified clusters, while individual maps show the distribution of CTLS from each time point across these clusters. (D) Quantification of cluster frequency among CTLs across the time course. Bars show the relative abundance of each protein-defined cluster in CTLs analyzed alone or after 2 h, 4 h, or 16 h of antigenic engagement. Cluster colors correspond to those shown in panel C. (E) Heatmap showing mean normalized marker expression across protein-defined CTL clusters. Markers are grouped according to early activation, cytotoxicity, cytokines, vesicular/degranulation and packaging/trafficking, late activation, and inhibitory receptor-associated categories. Data are from high-dimensional flow cytometry profiling of CMV-specific clonal CTLs. Expression values in the heatmap are scaled for each marker across clusters. Data are from three independent experiments.

Unsupervised clustering further partitioned the phenotypic landscape into discrete protein-defined CTL populations (Fig. 4C). Quantification of cluster abundance revealed a progressive redistribution of CTLs during antigenic engagement, with clusters enriched in CTLs analyzed alone or at early time points decreasing in relative abundance, while clusters 5 and 6 became predominant after 16 h (Fig. 4D). These two populations displayed distinct but overlapping phenotypic profiles, characterized mainly by differential expression of cytokines, late activation-associated proteins, and inhibitory receptors, together with broader changes in cytotoxic and vesicular machinery (Fig. 4E). Thus, prolonged antigenic engagement was not associated with convergence toward a single uniform endpoint, but with the emergence of multiple late phenotypes distinguished by different combinations of functional protein features. These findings raised the possibility that preserved population-level killing could coexist with progressive phenotypic diversification, a relationship examined directly in subsequent functional experiments.

Together, these data show that sustained antigenic engagement is associated with the redistribution of CTLs across a multidimensional protein-expression landscape characterized by distinct combinations of proteins spanning multiple functional categories. Together with the transcriptional analyses, these findings indicate that sustained antigenic engagement remodels multiple functional dimensions of the CTL phenotype with distinct temporal patterns. We therefore next asked whether this evolving protein landscape could be organized along a phenotypic remodeling trajectory.

### CTL functional-marker coordination is dynamically remodeled during sustained antigenic engagement

Having established that antigenic engagement alters the abundance of cytotoxic, vesicular/degranulation, and activation-associated proteins, we next asked whether these markers changed independently or whether their coordination was also remodeled at the single-cell level. To address this, we performed pairwise Spearman correlation analysis using single-cell protein-expression values from CMV-specific clonal CTLs analyzed at baseline or after 2 h, 4 h, or 16 h of engagement with pp65 peptide-pulsed JY target cells.

Correlation matrices revealed that CTL functional markers were not organized in a fixed relationship over time. At baseline, multiple cytotoxic and activation-associated markers showed coordinated expression, consistent with a pre-existing organization of the cytotoxic protein landscape (Supplementary Fig. 1). After 2 h of antigenic engagement, the correlation structure was altered, with several marker relationships weakened or redistributed. At later time points, particularly after 4 h and 16 h, new patterns of marker coordination emerged, involving cytotoxic, vesicular/degranulation, activation, and differentiation-associated markers (Supplementary Fig. 1).

Network visualization of significant marker correlations further supported this dynamic reorganization. Rather than showing a uniform increase or decrease in all marker relationships, CTLs displayed time-dependent changes in network architecture, with early engagement associated with partial disruption of baseline marker coordination and later engagement associated with reorganization of selected cytotoxic and activation-associated relationships (Supplementary Fig. 1). This suggests that sustained engagement does not merely alter individual marker abundance, but changes how functional markers are coordinated within single CTLs.

Together, these analyses indicate that CTL remodeling during sustained engagement involves dynamic reorganization of marker coordination at the single-cell level. This provided the rationale to next ask whether the remodeled protein landscape could be organized into functional CTL phenotypes associated with sustained cytotoxic potential.

### Trajectory analysis organizes multidimensional CTL phenotypic remodeling during sustained antigenic engagement

To determine how the multidimensional protein changes identified during antigenic engagement were organized over time, we performed principal component analysis of high-dimensional flow-cytometry profiles from CMV-specific clonal CTLs analyzed alone or after 2 h, 4 h, or 16 h of conjugation with pp65 peptide-pulsed JY target cells. We selected representative markers spanning activation and regulatory receptors, cytotoxic cargo, cytokines, and vesicular or degranulation-associated machinery to integrate these different functional dimensions within a common phenotypic framework. This analysis was intended to examine how their combined variation was organized during sustained antigenic engagement rather than to define complete protein programs.

Analysis of the integrated phenotypic marker panel separated CTLs along a principal-component axis associated with the duration of antigenic engagement. CTLs analyzed alone and at early time points were enriched at lower pseudotime values, whereas CTLs engaged for 16 h preferentially occupied later positions. Marker loadings indicated that progression along this axis reflected combined variation across several functional categories rather than changes in a single protein feature. Thus, sustained antigenic engagement was associated with progressive remodeling of the broader CTL protein phenotype (Fig. 5A-C).

**Figure 5.**
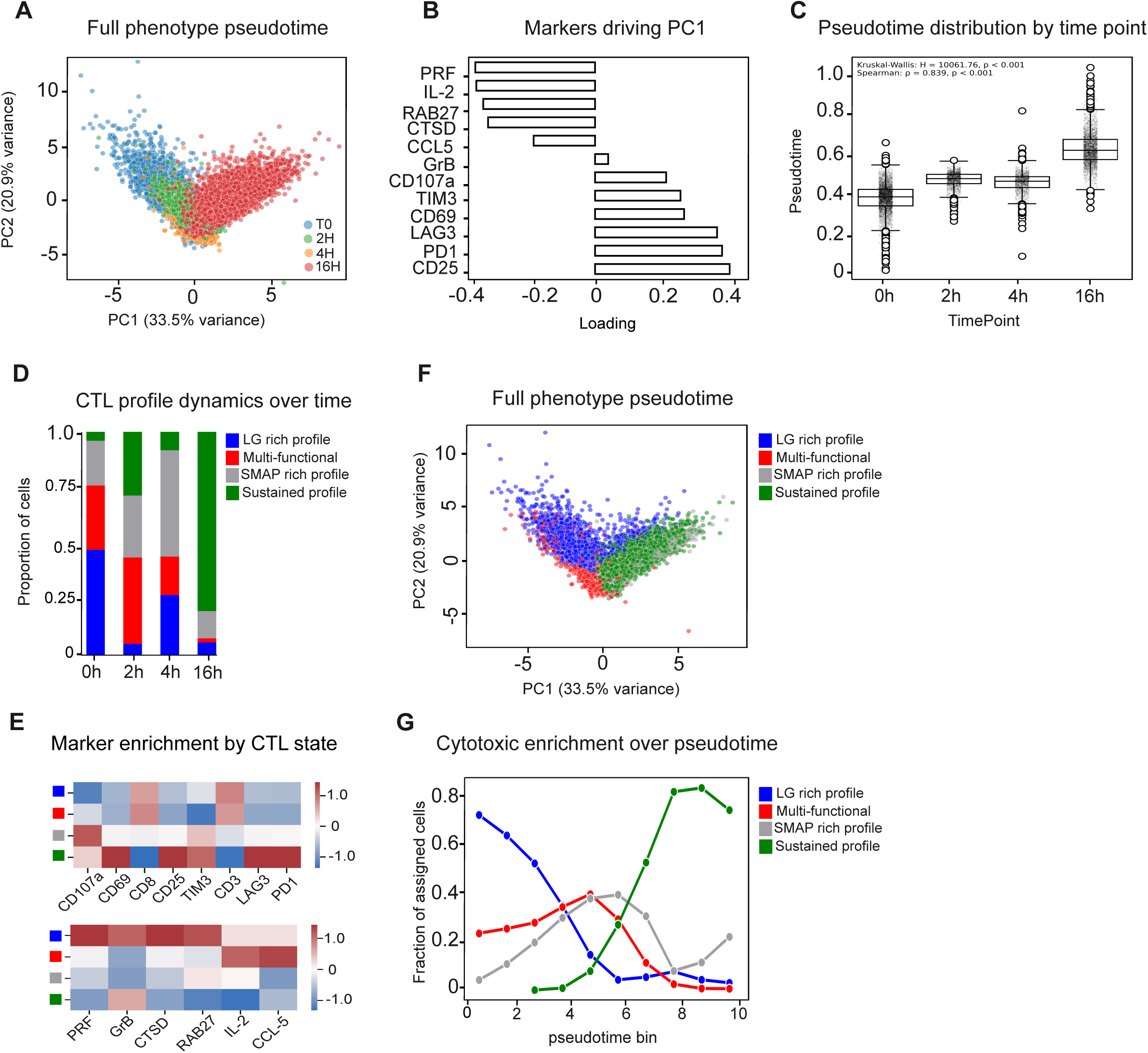
Trajectory analysis identifies CTL states associated with sustained cytotoxic potential. (A-C) Principal component analysis of high-dimensional flow cytometry profiles from CMV-specific clonal CTLs analyzed alone or after 2 h, 4 h, or 16 h of antigenic target-cell engagement. Cells are projected in PC1-PC2 space and colored according to experimental time point. (A) panel shows the PCA projection, (B) panel shows marker loadings contributing to PC1, and (C) panel shows pseudotime distribution across experimental time points. Pseudotime ordering was significantly associated with experimental time, with 16 h CTLs enriched at later pseudotime values. Each dot represents a single CTL and is colored according to experimental time point. (D) Relative abundance of CTL functional phenotypes across the time course. CTLs were classified into lytic-granule-rich, multifunctional, SMAP-like/secretory-remodeled, and sustained-like cytotoxic phenotypes based on cytotoxic and cytokine-associated marker composition. (E) Heatmaps showing marker enrichment across CTL functional phenotypes, including surface activation and inhibitory receptor markers, as well as intracellular cytotoxic and vesicular/degranulation-associated markers. (F) Projection of the defined CTL functional phenotypes onto the PCA-defined phenotypic landscape, showing that these phenotypes occupy distinct regions of the remodeling axis. (G) Relative abundance of each CTL functional phenotype across pseudotime bins, showing enrichment of lytic-granule-rich cells at early pseudotime and accumulation of SMAP-like/secretory-remodeled and sustained-like phenotypes at later pseudotime. Pseudotime was inferred using PCA-based ordering from either the full phenotypic marker panel or the cytotoxic machinery marker panel, as indicated. Data are representative of three independent experiments. Statistical significance was assessed using Kruskal-Wallis test for pseudotime differences across experimental time points and Spearman correlation for associations between pseudotime values, where indicated.

To resolve the phenotypic configurations underlying this remodeling, CTLs were classified according to combinations of cytotoxic-, cytokine-, vesicular-, and secretory-associated markers. This analysis identified lytic-granule-rich, multifunctional, SMAP-like/secretory-remodeled, and sustained-like cytotoxic phenotypes whose relative abundance changed over the time course (Fig. 5D). These phenotypes were defined by distinct combinations of protein features rather than by the expression of any single canonical cytotoxic marker.

Analysis of marker enrichment further demonstrated that the defined phenotypes differed across surface activation and inhibitory receptors, cytotoxic cargo, and vesicular or degranulation-associated proteins (Fig. 5E). Projection of these populations onto the PCA-defined phenotypic landscape showed that they occupied partially distinct regions along the remodeling axis (Fig. 5F). Their distribution along pseudotime was also temporally structured: lytic-granule-rich CTLs were enriched at early pseudotime, whereas SMAP-like/secretory-remodeled and sustained-like phenotypes accumulated at later pseudotime values (Fig. 5G).

Together, these data indicate that sustained antigenic engagement organizes CTLs along a multidimensional phenotypic remodeling axis, in which early lytic-granule-rich profiles give way to multiple later phenotypes characterized by different combinations of cytotoxic, secretory, activation, and regulatory protein features. Thus, prolonged engagement does not simply shift CTLs uniformly from one protein state to another, but progressively changes the relative representation of distinct multidimensional phenotypes.

### Surface-defined late-engaged CTL phenotypes display distinct cytotoxic capacity upon rechallenge

Having organized CTL protein remodeling into phenotypically distinct populations, we next asked whether late-engaged CTL phenotypes were associated with different functional capacities. High-dimensional analysis of CTLs after 16 h of antigenic engagement identified multiple phenotypically distinct populations within the late-engaged compartment (Fig. 6A). These populations were present at different relative abundances and displayed distinct marker-expression profiles, including differences in activation-associated surface proteins, cytotoxic machinery, and adhesion- and inhibitory receptor-associated features (Fig. 6B,C). Thus, CTLs exposed to the same duration of antigenic engagement remained phenotypically heterogeneous rather than converging on a uniform late state.

**Figure 6.**
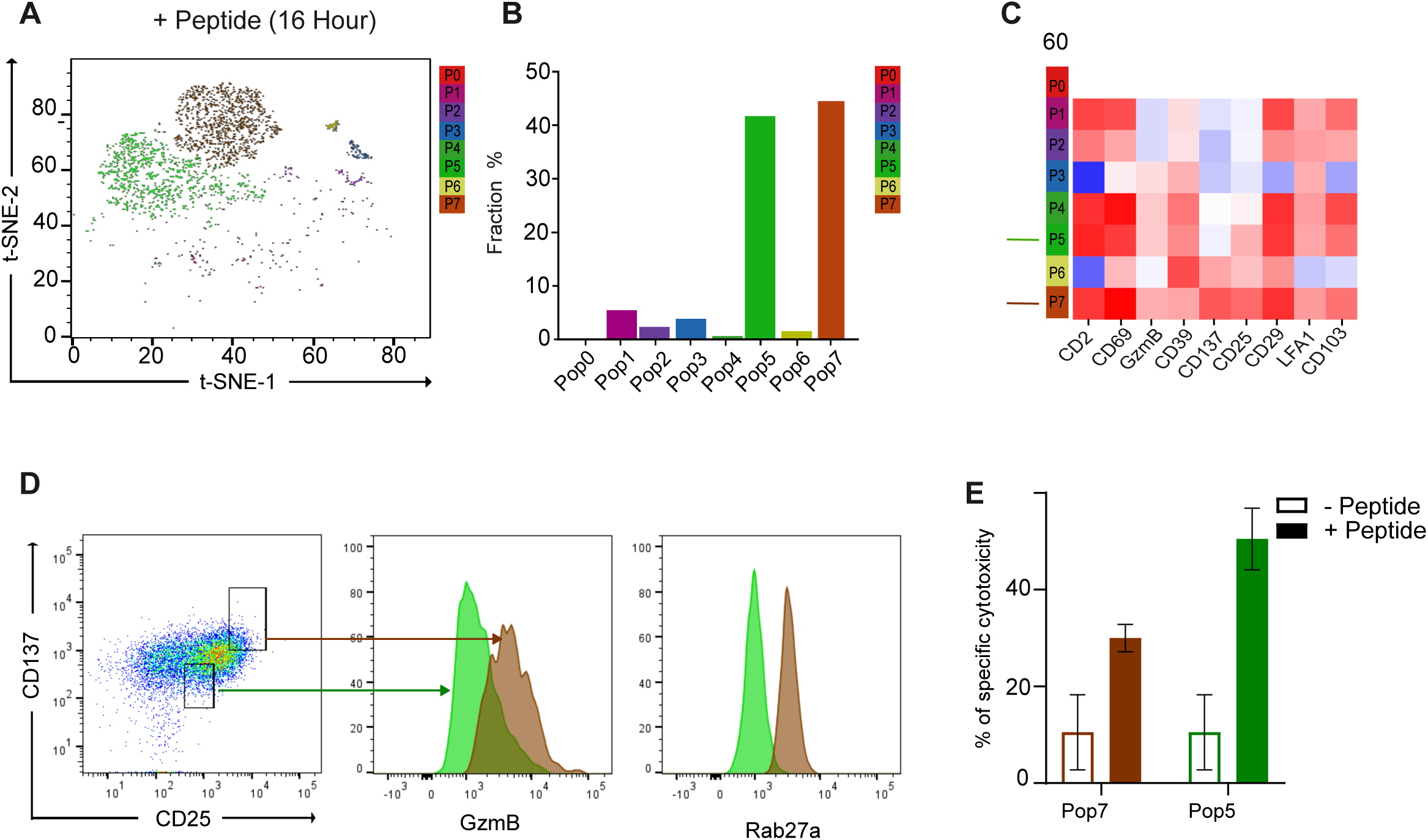
Surface-defined late CTL populations display distinct cytotoxic capacity upon rechallenge. (A) t-SNE projection of CTLs analyzed after 16 h of antigenic engagement, showing phenotypically distinct CTL populations identified within the late-engaged compartment. (B) Bar plot showing the relative abundance of each 16 h CTL population. (C) Heatmap showing marker-expression profiles of the indicated 16 h CTL populations, including CD25, CD69, CD137, cytotoxic markers, and adhesion/inhibitory receptor-associated markers. Pop5 and Pop7, selected for functional validation, are indicated. (D) Experimental strategy and sorting scheme used to isolate late CTL populations for functional validation. CTLs were first conjugated with pp65 peptide-pulsed JY target cells for 16 h and then subjected to FACS-based sorting according to CD25 and CD137 expression. CD25/CD137-defined populations corresponding to Pop5 and Pop7 were isolated. Histograms show GzmB and Rab27a expression in the sorted populations. (E) Rechallenge killing assay comparing the cytotoxic capacity of selected 16 h CTL populations against fresh unpulsed or pp65 peptide-pulsed target cells. Target-cell death was quantified after rechallenge to assess the functional capacity of the sorted CTL populations. Data are from high-dimensional flow cytometry profiling and functional rechallenge assays using CMV-specific clonal CTLs. Data are shown as mean +/-SEM from three independent experiments.

We selected Pop5 and Pop7 for functional analysis because they represented the predominant populations after 16 h of antigenic engagement. These populations could be distinguished using the surface markers CD25 and CD137 and differed in broader cytotoxic and vesicular-trafficking features (Fig. 6C). To isolate these populations without relying on intracellular markers, CTLs were conjugated with pp65 peptide-pulsed JY target cells and sorted after 16 h according to CD25 and CD137 expression (Fig. 6D). The resulting surface-defined populations showed differential GzmB and Rab27a expression, supporting their correspondence to phenotypes with distinct cytotoxic-cargo and trafficking characteristics.

The sorted CTL populations were subsequently rechallenged with fresh unpulsed or pp65 peptide-pulsed target cells. Rechallenge assays showed that the two late-engaged populations were not functionally equivalent, but displayed distinct capacities to induce death of fresh antigen-bearing target cells (Fig. 6E). These findings demonstrate that phenotypic heterogeneity within the late-engaged CTL compartment is associated with measurable differences in subsequent cytotoxic function.

Together, these data provide functional validation of the multidimensional phenotypes identified by high-dimensional cytometry. Although CD25 and CD137 were used as practical surface markers for population isolation, the sorted populations were originally defined by broader combinations of activation, cytotoxic, trafficking, adhesion, and regulatory protein features. Thus, their different killing capacities cannot be inferred from either CD25 or CD137 expression alone, but are associated with the multidimensional phenotypic configurations represented by these surface-defined populations. More broadly, these findings indicate that the maintenance of cytotoxic activity at the population level does not require all antigen-experienced CTLs to remain functionally equivalent. Early engagement is associated predominantly with lytic-granule-rich phenotypes, whereas sustained engagement is accompanied by diversification into phenotypically and functionally distinct CTL populations. The different cytotoxic capacities of these late populations are consistent with a model in which preserved bulk killing reflects the collective activity of a heterogeneous CTL compartment rather than the maintenance of a uniform fast-killing state.

### CAR T cells acquire features of the sustained cytotoxic remodeling state during repeated tumor-cell engagement

Having identified CTL phenotypic profiles associated with sustained cytotoxic function, we next asked whether elements of the same remodeling framework could be observed in therapeutic T cells. We therefore analyzed CAR T cells during repeated engagement with CD19^+^ target cells, first testing whether they retained cytotoxic capacity upon rechallenge and then asking whether this function was accompanied by acquisition of related phenotypic remodeling features.

To assess CAR T-cell cytotoxic capacity after prior tumor-cell encounter, CAR T cells were first cultured with Nalm-6 target cells and then rechallenged with a fresh target-cell population. CAR T cells efficiently killed the initially encountered target cells and retained cytotoxic activity against rechallenge targets after prior tumor-cell engagement (Fig. 7A). This indicated that, similar to the CTL model, CAR T cells retained cytotoxic activity following an initial phase of target-cell engagement under these assay conditions.

**Figure 7.**
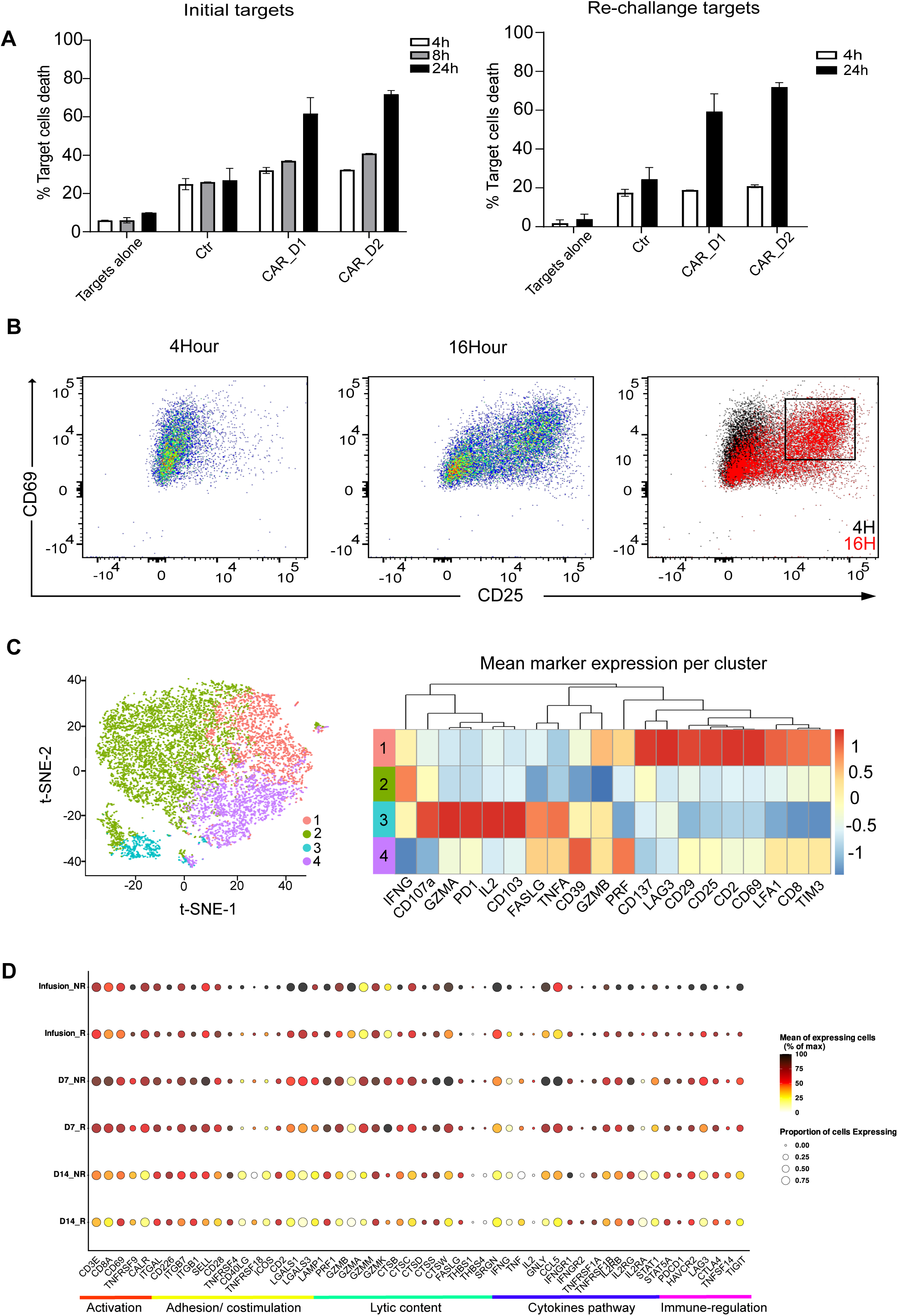
CAR T cells acquire features of multidimensional cytotoxic remodeling during repeated tumor-cell engagement. (A) Rechallenge killing assay assessing CAR T-cell cytotoxicity after initial exposure to Nalm6 target cells. CAR T cells were cultured with an initial population of target cells for the indicated duration and subsequently rechallenged with fresh target cells. Target-cell death was quantified in the initial and rechallenge target populations. Results are from two independent experiments performed in duplicates. (B) Flow-cytometric analysis of CD25 and CD69 expression in CAR T cells conjugated with RL lymphoma target cells for 4 h or 16 h. Representative plots show the redistribution of CAR T cells toward a CD25^high^CD69^high^ phenotype after prolonged target-cell engagement. Results are representative of two independent experiments, each performed in duplicate. (C) t-SNE projection and heatmap of protein expression showing the distribution of CAR T cells across multidimensional phenotypic states after 16 h of conjugation with RL target cells. (D) Longitudinal single-cell transcriptomic analysis of CD8 CAR T cells from tisa-cel-treated patients stratified according to clinical response. Dot plot shows expression of predefined genes representing functional categories derived from the CTL remodeling analyses in responder (R) and non-responder (NR) samples at infusion, day 7 (D7), and day 14 (D14) after infusion. Genes are grouped according to activation, adhesion/ contact interaction, cytotoxic machinery, cytokine-signaling, and regulatory/inhibitory receptor-associated programs. Dot size indicates the proportion of cells expressing each gene, and color indicates the mean expression among expressing cells, normalized to the maximum value for each gene. The day-14 analysis is shown descriptively because of the limited number of patients available at this time point. Marker categories are indicated below the plot.

We next asked whether this retained cytotoxic activity was accompanied by multidimensional phenotypic remodeling. Flow-cytometric analysis of CAR T cells conjugated with RL target cells showed an increase in CD25 and CD69 expression between 4 h and 16 h of tumor-cell engagement, with accumulation of a CD25^high^CD69^high^ population at the later time point (Fig. 7B). High-dimensional cytometry further showed that CAR T cells were organized into distinct protein-defined clusters during in vitro target-cell engagement (Fig. 7C). These clusters differed in the expression of activation, cytotoxic, inhibitory receptor, adhesion/ contact interaction, and cytokine-associated markers, indicating that repeated tumor-cell engagement redistributes CAR T cells across multidimensional protein phenotypes. Several of these features overlapped with those observed during sustained antigenic engagement of clonal CTLs, although the two cellular systems were not phenotypically identical.

Finally, we asked whether related functional dimensions were also evident in patient-derived CAR T cells and dynamically reorganized following infusion. We reanalyzed published single-cell RNA-sequencing data from tisa-cel-treated patients, restricting the analysis to CD8 CAR T cells and examining a predefined set of genes derived from our CTL remodeling analyses. These genes represented activation, adhesion/ contact interaction, cytotoxic machinery, cytokine signaling, and regulatory/inhibitory receptor-associated programs. Rather than seeking an identical transcriptional state across the experimental and clinical systems, we asked whether the organization of these functional features differed between clinical response groups and changed following in vivo CAR T-cell activation.

At infusion, responder and non-responder samples showed distinct distributions across the CAR T-cell transcriptional landscape, with responder cells enriched predominantly in clusters 0 and 1 and non-responder cells enriched in cluster 2 (Supplementary Fig. 2). At the product level, tisa-cel samples were largely concentrated within clusters 0-2, whereas axi-cel samples showed a broader distribution across transcriptionally defined populations, with greater representation of additional clusters (Supplementary Fig. 2C-D). Comparison of cluster transcriptional profiles indicated that these populations differed particularly in adhesion/contact interaction-associated features, which were more strongly represented in cluster 2. Consistent with this observation, the predefined remodeling-gene analysis showed the largest responder/non-responder differences in adhesion/contact interaction-associated genes at infusion, with more subtle differences persisting at day 7 (Fig. 7D).

By day 7, the features distinguishing responder and non-responder CAR T cells had shifted, with prominent differences in effector-associated genes including CCL5 and GNLY, together with residual differences in selected adhesion-associated genes such as ITGB1 (Fig. 7D). Thus, the molecular distinction between clinical response groups was not defined by a fixed marker or transcriptional program, but changed between the infusion product and the post-infusion state. Analysis of axi-cel-treated patients showed a less clearly organized response-associated transcriptional pattern across functional categories observed in the tisa-cel cohort (Supplementary Fig. 3).

Together, these findings indicate that CAR T cells retain cytotoxic competence during repeated target-cell engagement while undergoing multidimensional phenotypic remodeling. Patient-derived CAR T cells similarly occupied distinct combinations of functional transcriptional programs, and the features distinguishing responder and non-responder tisa-cel samples changed between infusion and day 7. Rather than identifying a single conserved cytotoxic state, these data support the extension of the CTL remodeling framework to therapeutic T cells as a dynamic reorganization of multiple functional dimensions during continued antigenic challenge.

## Discussion

How CTLs preserve cytotoxic function during sustained antigenic engagement remains less well understood than the mechanisms governing an individual killing event. Our data reveal that this persistence is not achieved by maintaining an unchanged cytotoxic state. Antigen-experienced CTLs retained overall cytotoxic activity upon rechallenge, yet the targets they engaged displayed altered death dynamics. PI entry was delayed and reduced in intensity, the intermediate PI-positive population was markedly decreased, and extensive membrane blebbing developed more slowly, while the overall fraction of targets eliminated during the 2 h assay remained comparable. Thus, preserved population-level killing did not imply preservation of the same cytotoxic behavior or cellular configuration. This dissociation provided the starting point for examining how CTLs remodel while maintaining cytotoxic competence during sustained antigenic engagement.

The first consequence concerns how cytotoxic capacity is estimated, which is routinely inferred from GzmB, IFNγ, CD107a or selected activation receptors, although target-cell killing depends on the coordinated contribution of cytotoxic cargo, vesicular and degranulation machinery, adhesion and contact-interaction molecules, cytokines, and regulatory receptors. Previous studies have shown that lytic and stimulatory components of CTL function can be uncoupled and that clonally related CTLs sharing antigen specificity can nevertheless differ substantially in killing efficiency, serial cytotoxicity, and cytokine production(Cazaux et al., 2019; Faroudi et al., 2003; Khazen et al., 2021; Purbhoo et al., 2004; Varadarajan et al., 2012; Vasconcelos et al., 2015). Our findings extend this concept by showing that late antigen-engaged CTLs occupy multidimensional phenotypes with distinct subsequent killing capacities. CD25 and CD137 provided practical surface markers for isolating Pop5 and Pop7, but these populations were originally distinguished by broader combinations of activation, cytotoxic, trafficking, adhesion, and regulatory features. Their functional difference should therefore be interpreted in the context of these multidimensional phenotypes rather than attributed to either surface marker alone.

The altered target-cell death dynamics observed after prior CTL engagement further raise the possibility that sustained cytotoxicity involves changes not only in the magnitude of killing but also in how killing is executed. Reduced PI uptake and delayed membrane blebbing are consistent with altered membrane-damage kinetics and may reflect changes in the relative contribution, timing, or efficiency of the cytotoxic mechanisms used during rechallenge. Several mechanisms can contribute to CTL-mediated target-cell death, including perforin- and granzyme-dependent lysis, death-receptor pathways, and the delivery of supramolecular attack particles (Bálint et al., 2020; McKenzie et al., 2022). The present experiments do not distinguish their relative contributions, and direct perturbation of individual cytotoxic pathways would be required to determine whether prior engagement modifies killing modality. Nevertheless, the dissociation between altered target-cell death kinetics and preserved overall killing indicates that equivalent cytotoxic output can arise from functionally distinct cytotoxic configurations.

The time-resolved transcriptomic analysis further showed that the functional components of the CTL response did not remodel synchronously. Cytotoxic effector- and cytokine-associated programs increased rapidly during early antigenic engagement, together with activation-related programs, whereas vesicular-trafficking and granule-packaging programs followed distinct intermediate kinetics. In contrast, adhesion/contact interaction-associated programs progressively declined during sustained engagement. Regulatory and inhibitory receptor-associated programs also evolved dynamically across the time course. Sustained antigenic stimulation therefore did not simply amplify a pre-existing effector program but progressively reorganized different functional dimensions with distinct kinetics. The delayed remodeling of vesicular and granule-associated programs may reflect the requirement to replenish or reorganize secretory machinery following early cytotoxic activation, although the mechanisms controlling this temporal organization remain to be established. More broadly, these data indicate that estimates of cytotoxic potential are inherently dependent on both the functional dimension measured and the time at which it is examined.

Regulatory and inhibitory receptor features appeared during the same window while CTLs remained able to kill. Sixteen hours is far shorter than the timescales over which chronic stimulation produces stable functional impairment, and inhibitory receptor expression alone has been argued to be insufficient to define exhaustion (Blank et al., 2019), with activation-driven expression separable from the developmental states that characterize exhausted CD8 T cells(Beltra et al., 2020). In our system these proteins therefore index recent and ongoing TCR signalling rather than a differentiation endpoint. The distinction matters for the interpretation of early phenotyping data, where detection of PD-1, TIM-3 or LAG-3 is frequently taken as evidence that effector function has been compromised.

Trajectory analysis ordered the protein-expression landscape from early lytic-granule-rich profiles toward several later phenotypic configurations, and functional isolation demonstrated that two late populations were not equivalent in their ability to kill fresh targets. CTLs exposed to the same antigenic environment for the same overall duration therefore remained phenotypically and functionally heterogeneous. Equal time in culture, however, does not imply equivalent engagement history. Individual CTLs are likely to differ in the number and duration of target-cell encounters experienced during the assay, and the late phenotypic diversity observed here may therefore reflect differences in cumulative engagement history as well as pre-existing or engagement-induced cellular heterogeneity. Distinguishing these possibilities will require experimental approaches that link the encounter or degranulation history of individual cells to their subsequent phenotype and function. What the present data establish is that preserved population-level cytotoxicity can coexist with substantial divergence in the phenotype and subsequent killing capacity of the cells composing that population, without implying that the identified populations directly cooperate or compensate for one another.

The phenotypic classifications used here should therefore be considered operational descriptions of a continuous remodeling landscape rather than fixed CTL identities. Their purpose is to organize recurrent combinations of functional features and to identify populations that can subsequently be tested experimentally. In particular, the designation of the secretory-remodeled population reflects its combination of trafficking- and cargo-associated features rather than direct demonstration of supramolecular attack-particle formation. Direct visualization or molecular characterization of these structures would be required to establish such an interpretation.

The extension of this analysis to CAR T cells suggests that multidimensional remodeling is not restricted to conventional TCR-mediated cytotoxicity. CAR T cells retained cytotoxic activity following prior target-cell encounter and redistributed across protein-defined phenotypes during prolonged tumor-cell engagement. These phenotypes shared several functional dimensions with those identified in clonal CTLs, including variation in activation, cytotoxic, adhesion/ contact interaction, cytokine, and inhibitory receptor-associated features, but the two systems were not phenotypically identical. This partial overlap is consistent with the distinct signaling architecture and synaptic organization of CAR- and TCR-mediated activation(Davenport et al., 2018; Salter et al., 2018). Thus, the relevant level of similarity between these systems may lie in the dynamic reorganization of related functional axes rather than in convergence toward an identical cellular state.

Analysis of patient-derived CD8 CAR T cells further indicated that these functional dimensions remain dynamically organized *in vivo*. In tisa-cel-treated patients, responder and non-responder samples differed predominantly in adhesion/ contact interaction-associated features within the infusion product, whereas by day 7 the distinction involved more prominent differences in effector-associated genes including CCL5 and GNLY. Previous studies have established that the cellular and transcriptional composition of the infusion product is associated with clinical outcome and that CAR T-cell products differ substantially in their cellular composition (Deng et al., 2020; Fraietta et al., 2018; Haradhvala et al., 2022). Our analysis therefore does not identify a new response-associated state, but instead highlights that the functional features distinguishing clinical response groups are themselves time dependent.

Several limitations constrain the interpretation of these clinical observations. The number and timing of antigenic encounters experienced by CAR T cells in patients cannot be reconstructed from the available samples, and post-infusion transcriptional changes therefore cannot be attributed directly to repeated tumor-cell killing. The limited number of samples at later time points also prevents association of particular remodeling trajectories with clinical outcome. Moreover, projecting functional programs identified in an in vitro clonal system onto patient-derived CAR T cells establishes overlap in the biological dimensions being remodeled, but does not demonstrate that the same underlying process drives these changes in vivo. The greater inter-patient heterogeneity observed in the axi-cel cohort may similarly reflect differences in CAR design, product composition, patient characteristics, or other variables that cannot be resolved in the present dataset.

Taken together, these findings support a view of sustained cytotoxicity as a dynamic population property rather than the persistence of a single optimal effector state. Population-level killing can remain stable while the CTLs producing that output progressively reorganize across cytotoxic, secretory, activation, interaction, and regulatory dimensions and acquire different capacities to respond to subsequent target encounters. Consequently, bulk cytotoxicity measurements or individual effector markers may conceal substantial changes in the functional composition of an antigen-experienced T-cell population. Characterizing sustained cytotoxic function may therefore require resolving how multidimensional cellular configurations evolve over time, in addition to measuring the magnitude of cytotoxic output itself.

## Acknowledgements

We thank Dr. Capucine Grandjean for helpful discussions and critical input. We thank the flow cytometry core facilities of the Centre de Recherches en Cancérologie de Toulouse (CRCT), especially Manon Farcé for technical support. Part of flow cytometry experiments were performed at the Univ Toulouse, INSERM, CNRS, Infinity core facility connected to Toulouse Réseau Imagerie network, member of the France-BioImaging national infrastructure supported by the French National Research Agency (ANR-24-INBS-0005 FBI BIOGEN). We thank Anne-Laure Iscache for the standardization of the flowcytometry panel, training personnel and development of analytic pipeline. We also thank Fatima l’faqihi for her expert advice on the set up. This work was supported by the Fondation ARC pour la recherche sur le cancer, and an IUCT-Oncopole Translational Research Grant secured by R.K. and by the European Research Council (ERC) under the European Union’s Horizon 2020 Research and Innovation Programme (Grant agreement No. Syn-951329) secured by S.V. The funders had no role in study design, data collection and interpretation, or preparation of the manuscript.

## Author contributions

R.K. conceived and led the study, designed the project, supervised the research, analyzed and interpreted data, secured funding and wrote the paper. A.J. contributed to the design of the research and led the analysis and interpretation of the single-cell transcriptomic data. H.R.G. performed the imaging and flow-cytometry experiments and analyzed data. F.A., M.E. and M.T. analyzed single cell RNA seq data, and F.A. prepared the figures. A.L.I. developed the flow-cytometry panel and analysis pipeline. S.M. and B.C. generated the CTL clones and CAR T cells, respectively. A.V. performed CAR T cell experiments and analyzed data. C.L. provided clinical expertise and discussed results. S.V. provided financial support, discussed results and edited the paper. All authors reviewed and approved the paper. The authors declare no competing interests.

## Material and Methods

### Cellular model

HLA-A2-restricted CD8^+^ T cells specific for the NLVPMVATV peptide of the cytomegalovirus protein pp65 (CMV specific) clones already available in the lab were used in this study. Cells were cultured in RPMI 1640 medium supplemented with 5% human AB serum (Inst. Biotechnologies J.BOY), 50µmol/L 2-mercaptoethanol, 10mM HEPES, 1X MEM NEAA (Gibco), 1X Sodium pyruvate (Sigma), 10µg/ml ciprofloxacine (AppliChem), 100 IU/ml human rIL-2 and 50 ng/ml human rIL-15 (Miltenyi). T cell clones were stimulated in complete RPMI/HS medium containing 1 µg/ml PHA with 1 x 10^6^ per ml 35 Gy irradiated allogeneic peripheral blood mononuclear cells (isolated on FicollPaque Gradient from blood of healthy donors). Re-stimulation of clones were performed every 2 weeks. Blood samples from healthy donors were collected and processed following standard ethical procedures, after obtaining written informed consent from each donor and approval from the French Ministry of Research (transfer agreement AC-2020-3971). Authorization for the preparation and conservation of cell lines and clones derived from healthy donor blood samples was granted by the French Ministry of Research (authorization no. DC-2021-4673).

As target cells, EBV-transformed B cells (JY) HLA-A2^+^ were used with high sensitivity to CTL attack that are routinely used in our lab to study CTL behavior. RL cells, a human B-cell lymphoma cell line derived from a patient with follicular lymphoma, and NALM-6 cells, a human B-cell acute lymphoblastic leukemia (B-ALL) cell line, were used as targets in CAR T-cell experiments. Target cells are cultured in RPMI 1640 GlutaMAX supplemented with 10% FCS and 10mM HEPES, 1X MEM NEAA (Gibco), 1X Sodium pyruvate (Sigma), 10µg/ml ciprofloxacine (AppliChem).

All cell lines were screened biweekly for mycoplasma contamination using MycoAlert mycoplasma detection kit (Lonza).

### CTL/Target cell conjugation

Target cells were left unpulsed or pulsed with indicated concentration of antigenic peptide (pp65: NLVPMVATV) during 2h at 37°C/5% CO2, washed three time, re-suspended in RPMI 5% FCS/HEPES (R5). CTLs were pre-stained with 0.1µM CMFDA (5-chloromethylfluorescein diacetate (Molecular Probes)), or CTV (Celltrace Violet (Molecular Probes)) for 15min at 37°C/5% CO2, washed, re-suspended in R5 and added to the target cells at different CTL/target cell (E:T) ratio. For flow cytometry-based read-outs, cells were seeded to a 96-wells U-bottom plate and pelleted for 1min, 1500rpm and subsequently incubated at 37°C/5% CO2 for the indicated time point. For imaging-based analysis, CTL/target cells seeding were performed either in 15-well chambered slides (Ibidi, Munich, Germany) or in slides.

### Live cell imaging of conjugates

Target cells preloaded with 10µM of antigenic peptide were seeded at 1.5 × 10^5^ cells per well on poly-D-lysine-coated 15-well chambered slides 5 min before imaging. Chambered slides were mounted on a heated stage within a temperature-controlled chamber maintained at 37 °C, and constant CO_2_ concentrations (5%). To monitor cytotoxic modality CTLs and target cells were conjugated at the E:T ratio of 1:1. Effector CTLs were either fresh or previously conjugated with targets, and were added at the start of recording. CTLs were optionally labeled with 1 μM Cell Tracer Violet (Invitrogen) for visualization. To monitor membrane permeabilization, propidium iodide (PI; 0.5 μg/mL, Sigma) was included in the medium. A rise in PI fluorescence was interpreted as a marker of membrane disruption, often associated with perforin pore formation. Images of CTL/target cell conjugates were acquired for 2 h using either a Nikon spinning disk microscope (60×/1.4 NA Plan Apochromat LBDA objective lens (Nikon Instruments)). Image acquisition was controlled using MetaMorph Software (Version 7.10.5.476, Molecular Devices) and Modular V2.0 GATACA software. A sCMOS Hamamatsu ORCA-Flash4.0 V3 camera or an emCCD Evolve (Photometrics) camera were used for acquisitions. Target cells were preloaded with 5 ng/µl Fluo-4 AM, (Molecular Probes) during 30 min and washed before seeding in chambered slides. Image analysis was performed using Image J software.

### CTL cytotoxicity assay

CTL/target cell conjugates were transferred on ice and disrupted by several rounds of pipetting. Before reading samples on flow-cytometry, 0.25 μg of 7-aminoactinomycin D (7-AAD; BD Biosciences) was added to each sample to measure targets death.

### Single-cell RNA sequencing

Following coculture with target cells for the indicated durations, viable cells were sorted using a BD FACSAria cell sorter. Dead cells were excluded based on 7-AAD staining, and a forward- and side-scatter gate corresponding to the size and granularity of CTLs was applied to enrich the collected fraction for CTLs. Because no CTL-specific surface marker was used during sorting, the resulting samples contained both CTLs and residual target cells, with CTLs enriched on the basis of their scatter characteristics. Sorted cells were collected in chilled complete culture medium and immediately processed for single-cell RNA sequencing.

For the transcriptomic analysis presented in figure 2, single-cell gene-expression libraries were generated using the Chromium Single Cell 5′ Gene Expression platform and the Chromium Next GEM Single Cell 5′ reagent kit, according to the manufacturer’s instructions (10x Genomics). T-cell receptor V(D)J libraries were not generated. For the experiment presented in Figure 3, libraries were generated using the Chromium Single Cell 3′ Gene Expression platform and the Chromium Next GEM Single Cell 3′ reagent kit. Eight samples were barcoded using the 10x Genomics CellPlex sample-multiplexing system and processed in two multiplexed pools of four samples each. These comprised CTLs cultured alone, with unpulsed JY target cells, or with pp65 peptide-pulsed JY cells for 2, 4, 6, or 16 h, together with two additional samples that are excluded from the analyses presented in this manuscript. Sample identity was recovered computationally from sample-specific oligonucleotide barcodes. Gene-expression libraries were sequenced on an Illumina NovaSeq system.

Raw sequencing data were processed using Cell Ranger software (10x Genomics). Reads were demultiplexed and aligned to the human GRCh38 reference genome using the 10x Genomics reference package refdata-gex-GRCh38-2020-A, and used to generate gene-barcode expression matrices. For the multiplexed experiment, sample-specific barcode information was processed using Cell Ranger multi to assign individual cells to their sample of origin and to identify and exclude unassigned cells and cross-sample multiplets. The resulting filtered feature-barcode matrices were imported into Seurat in R for quality control, normalization, dimensionality reduction, unsupervised clustering, differential-expression analysis, and downstream transcriptomic analyses. CTLs and target cells were subsequently distinguished according to their transcriptomic profiles and expression of lineage-specific markers.

### Functional phenotyping of CTL with flow-cytometry

Using the same experimental set up, CTL/target cell conjugates were stopped at different time points and subjected to staining for the expression of several markers involved in CTL activation and degranulation. CTL/target cells conjugates were washed in ice-cold PBS containing 1% human serum, 1% FBS and 0.5 mM EDTA. Non-specific bindings were prevented by 15 min incubation with PBS containing 1% FCS which is used throughout the procedure as staining and washing buffer. Cells were stained first extracellularly with a mix of specific antibodies (anti-CD3 and anti-CD107a) for 30 minutes on ice. Samples were washed thoroughly and fixed with 3% PFA, permeabilized with 0.1% saponin (in PBS/3% BSA/HEPES), and stained for intracellular markers (anti-perforin and anti-GranzymeB). Samples were acquired using Spectral Aurora 5A (Cytek) or Symphony A5 (BD Biosciences). Results are analyzed using the FlowJo Pro software (Tree Star, Inc.). The following mouse anti human antibodies were used: Alexa Fluor APC-Cy7 or Alexa Fluor 647 anti-perforin (5 μg/ml, clone dG9; BioLegend), BV421 anti-perforin (5 μg/ml, clone BD48; BioLegend), PercpCy5.5 or BUV395 anti-CD107a (10 μg/ml, clone H4A3; BD Biosciences), Pacific Blue anti-granzyme B (10 μg/ml, clone GB11; BioLegend) and PE anti-CD3 (10 μg/ml, clone OKT3; BD Biosciences), SuperBright783 anti-CD3 (10 μg/ml, clone SK7; BD Biosciences), BV570 anti-CD8 (10 μg/ml, clone RPA-T8; BD Biosciences), BV711 anti-CD69 (10 μg/ml, clone FN50; BD Biosciences), Alexa Fluor 594 anti-Cathepsin D (10 μg/ml, clone H.5; Santa Cruz), Alexa Fluor 680 anti-Rab27a (10 μg/ml, clone E.8; SantaCruz), RB744 anti-TNF (10 μg/ml, clone N297A; BD Biosciences), APC anti-IFNy (10 μg/ml, clone 4S.B3; BD Biosciences), BB700 anti-IL-2 (10 μg/ml, clone MQ1-17H12; BD Biosciences), PE-Fire700 anti-PD1 (10 μg/ml, clone A17188B; BioLegend), PE-Cy7 anti-LAG-3 (10 μg/ml, clone 11C3C65; BD Biosciences), SuperBright783 anti-TIM3 (10 μg/ml, clone Ber-ACT8; BD Biosciences). BUV395 anti-CD137 (10 μg/ml, clone 4B4-1; BD Biosciences), BV650 anti-CD25 (10 μg/ml, clone 2A3; BD Biosciences), PE-Cy7 anti Granzyme A (10 μg/ml, clone CB9; BioLegend), PE anti-FasL (10 μg/ml, clone NOK-1; BioLegend), Alexa Fluor 488 anti-Serglycin (10 μg/ml, clone C-11; SantaCruz), PE anti CCL (10 μg/ml, clone 2E9; BioLegend).

### High dimensional analysis

CTLs were identified as live CMFDA⁺ cells, and were exported as FCS files and analyzed using FlowJo v10 with the t-SNE and FlowSOM plugins. Before dimensionality reduction, fluorescence intensities were transformed using an arcsinh transformation with a cofactor of 150. Equal numbers of CTLs were randomly downsampled from each condition and concatenated to avoid bias due to unequal cell numbers. The markers used for dimensionality reduction and clustering included CD3, CD107a, GzmB, GzmA, perforin, FasL, IFNγ, TNFα, cathepsin D, serglycin, CD25, CD69, PD-1, TIM-3, LAG-3. Markers used only for gating, cell identification, or viability exclusion were not included in the t-SNE or FlowSOM computation. t-SNE analysis was performed on the concatenated CTL population using 2000 events per condition, with the following parameters: perplexity 30, iterations 1000, automatic learning rate, and theta 0.5. The resulting t-SNE maps were visualized according to time point, antigenic stimulation, target-cell type, or FlowSOM cluster assignment, as indicated in the corresponding figure legends. FlowSOM clustering was performed on the same transformed and downsampled CTL dataset using the selected phenotypic markers. The self-organizing map was generated using a 10 × 10 grid, followed by metaclustering into several phenotypic clusters. Cluster identity was assigned based on marker-expression profiles and visualized on the corresponding t-SNE maps. Cluster frequencies were calculated as the percentage of live CTLs within each condition. Median fluorescence intensity values for individual markers were extracted from manually gated CTLs or from FlowSOM-defined clusters, as indicated. For pooled quantifications, values were normalized to the corresponding non-pulsed condition or to the indicated baseline condition within each independent experiment before statistical analysis.

### Single-cell flow cytometry analysis of CTL cytotoxic-state remodeling

Single-cell flow cytometry data from CTLs analyzed at 0 h, 2 h, 4 h, and 16 h after antigenic engagement were imported from CSV files and merged into a single data frame prior to normalization. Cytotoxic-state scores were calculated using z-scored expression of Perforin, Granzyme B, Cathepsin D, RAB27, IL-2, and CCL5. Specifically, the lytic-granule-rich score combined Perforin, Granzyme B, and Cathepsin D expression; the sustained-like score reflected high Granzyme B relative to Perforin; the SMAP-like/secretory-remodeled score reflected high RAB27 together with low Perforin and Cathepsin D; and the multifunctional score combined IL-2 and CCL5 expression. Cells were assigned to the state with the highest score. Cells for which the difference between the highest and second-highest score was below 0.2 were classified as unassigned and excluded from state-enrichment analyses. Global phenotypic remodeling was assessed by PCA using cytotoxic, cytokine-associated, degranulation, activation, and inhibitory receptor markers. Pseudotime was inferred from the principal component most strongly correlated with experimental time, oriented to increase from 0 h to 16 h, and normalized to a 0–1 scale. Cytotoxic-state enrichment along pseudotime was determined by dividing cells into ten equal-frequency pseudotime bins and calculating the fraction of each assigned state per bin. Statistical differences in pseudotime distributions across experimental time points were assessed using Kruskal-Wallis tests. Analyses were performed in Python using pandas, NumPy, scikit-learn, SciPy, seaborn, and matplotlib.

### Human CD19-CAR T cells generation

Human CD8⁺ anti-CD19 CAR T cells were kindly provided by Philippe Bousso’s laboratory and generated as previously described by Boulch et al. Briefly, CD8⁺ T cells were isolated from peripheral blood mononuclear cells of healthy donors obtained from the Etablissement Français du Sang (EFS) and activated with anti-CD3/CD28 Dynabeads. Cells were subsequently transduced with a lentiviral vector encoding an anti-CD19 FMC63-based CAR containing a CD8 transmembrane domain and CD28-CD3ζ signaling domains. Following transduction and removal of the activation beads, CD8⁺ CAR T cells were used for in vitro cytotoxicity assays. Alternatively, the same CAR construct was transiently expressed in CD8⁺ T cells by electroporation of CAR-encoding mRNA obtained from Addgene, using the Lonza 4D-Nucleofector system with the P3 Primary Cell Nucleofector Kit, according to the manufacturer’s instructions. Electroporated CAR T cells were subsequently used for rechallenge cytotoxicity assays.

### Patient-derived CAR T-cell scRNA-seq analysis

Published single-cell RNA-sequencing data from patients treated with tisa-cel or axi-cel were reanalyzed from Haradhvala et al. (2022). Analyses were restricted to CD8 CAR T cells and performed separately for each CAR T-cell product. Responder and non-responder samples were compared at the infusion-product and day-7 time points; day-14 samples were analyzed descriptively where available.

Cells were analyzed using Seurat in R, including dimensionality reduction, unsupervised clustering, and UMAP visualization. Expression of predefined genes representing activation, adhesion/ contact interaction, cytotoxic machinery, cytokine signaling, and regulatory/inhibitory receptor-associated programs was compared between response groups at each time point. Because patient numbers were limited at later time points, these analyses were considered exploratory.

