## Supplementary-information for "Sustained antigenic engagement programs multidimensional CTL states with distinct cytotoxic capacities"

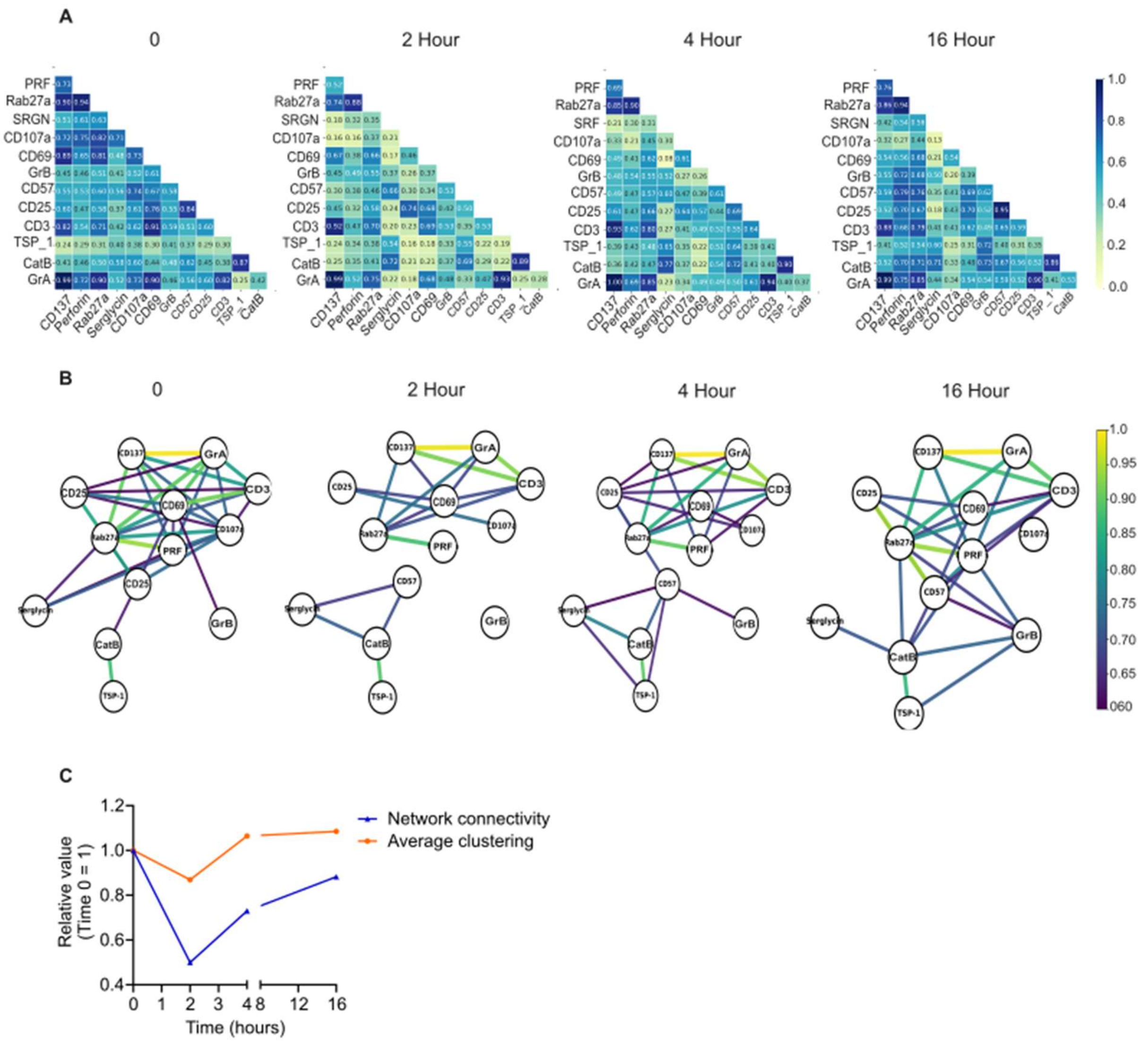


**Supplementary figure 1. CTL functional-marker coordination is dynamically remodeled during sustained engagement.**

(A) Spearman correlation matrices showing pairwise relationships between cytotoxic, vesicular/degranulation, activation, and differentiation-associated protein markers in CMV-specific clonal CTLs analyzed at baseline or after 2 h, 4 h, or 16 h of conjugation with pp65 peptide-pulsed JY target cells. Correlation coefficients are shown for each marker pair. (B) Network representation of marker correlations at each time point. Nodes represent individual protein markers and edges represent marker pairs passing the indicated correlation threshold. Edge thickness and color intensity indicate correlation strength. (C) Quantification of network properties across the time course Network connectivity was calculated as the number of threshold-passing positive correlations and normalized to baseline. Average clustering coefficient was used as a measure of local network organization and was also normalized to baseline. Values are shown relative to T0. Correlation analyses were performed using single-cell marker-expression values from high-dimensional flow cytometry. Networks were generated using a fixed correlation threshold of Spearman r ≥ 0.6.

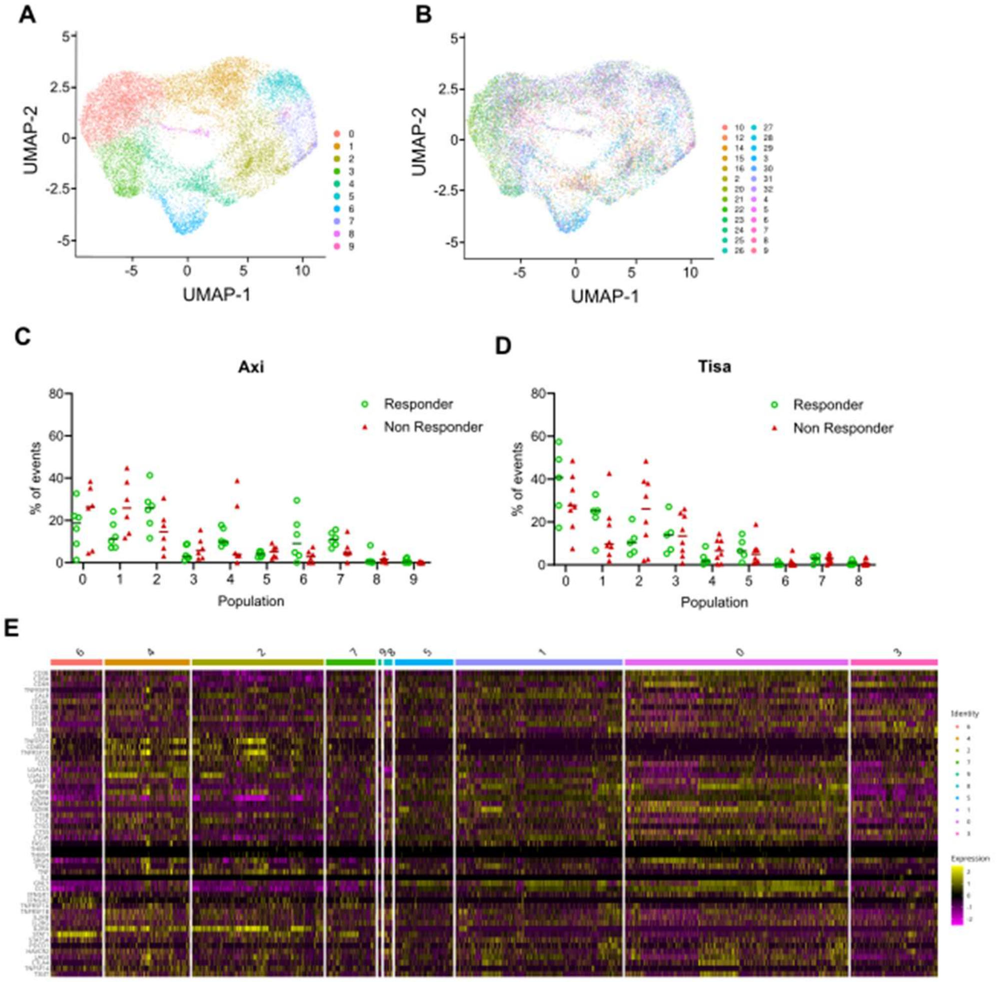


**Supplementary Figure 2. Transcriptional organization of CD8 CAR T cells in axi-cel- and tisa-cel-treated patients.**

(A) UMAP projection of CD8 CAR T cells colored according to unsupervised transcriptional cluster identity. (B) UMAP projection of the same cells colored according to individual patient identity. (C-D) Relative abundance of transcriptional clusters across individual patients treated with axi-cel (C) or tisa-cel (D). Each point represents one patient and indicates the proportion of CD8 CAR T cells assigned to the corresponding transcriptional cluster, illustrating inter-patient variation in cluster representation within each CAR T-cell product. (E) Heatmap showing expression of selected genes across transcriptional clusters. Genes are grouped according to predefined functional categories, including activation, adhesion/contact interaction, cytotoxicity, cytokine-associated, and regulatory/inhibitory receptor-associated programs. Cluster 2 showed stronger representation of adhesion/contact interaction-associated features.

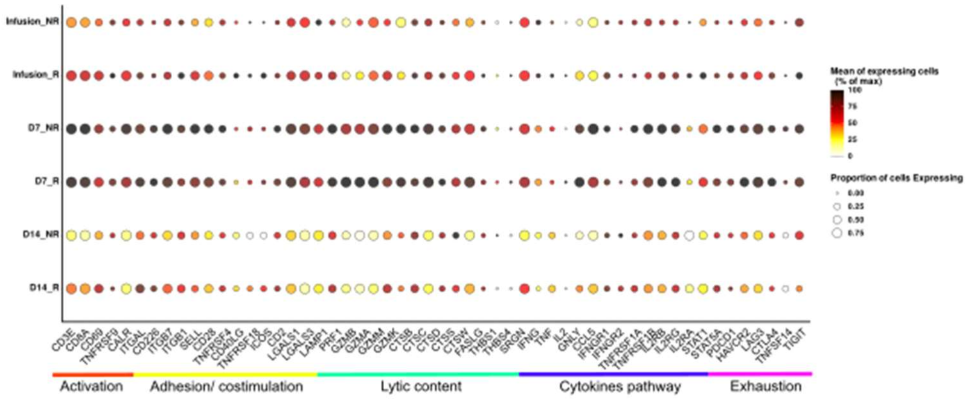


**Supplementary Figure 3. Functional transcriptional profiles of CD8 CAR T cells from axi-cel-treated patients.**

Dot plot showing expression of predefined genes representing activation, adhesion/contact interaction, cytotoxic machinery, cytokine signaling, and regulatory/inhibitory receptor-associated programs in CD8 CAR T cells from axi-cel-treated patients. Dot size indicates the proportion of cells expressing each gene, and color indicates mean expression among expressing cells, normalized to the maximum value for each gene.
